# AtMYB50 regulates root cell elongation by upregulating *PECTIN METHYLESTERASE INHIBITOR 8* in *Arabidopsis thaliana*

**DOI:** 10.1101/2023.04.19.537493

**Authors:** Kosuke Mase, Honomi Mizuno, Norihito Nakamichi, Takamasa Suzuki, Takaaki Kojima, Sho Kamiya, Taiga Takeuchi, Chiko Kondo, Harumi Yamashita, Satomi Sakaoka, Atsushi Morikami, Hironaka Tsukagoshi

## Abstract

Plant root development is regulated by several signal transduction pathways. Among them, plant phytohormones, like auxin and cytokinin, are well characterized for their molecular mechanisms of action. Reactive oxygen species (ROS) play important roles as signaling molecules in controlling root development. Under these signal transduction pathways, the gene regulatory network, which is controlled by transcription factors, is the key to regulating root growth. We have previously reported an important transcription factor, *UP BEAT1* (*UPB1*), that regulates ROS homeostasis at the root tip, further controlling the transition from cell proliferation to differentiation. Although UPB1 directly regulates the expression of several peroxidases that control ROS homeostasis, UPB1 still targets genes other than peroxidases. This indicates that UPB1 may regulate root growth through different ROS signals. Here, we investigated the function of the transcription factor *MYB50*, a direct target of UPB1, in *Arabidopsis thaliana*. We then examined whether UPB1 regulates *MYB50* expression in the roots using an induction expression system and imaging of multiple fluorescent proteins. We also performed RNA-Seq analysis using *MYB50* estradiol induction lines and ChIP-seq analysis to identify the *MYB50* regulatory gene network. Integrating these analyses with *UPB1* regulatory network revealed that MYB50 regulates the expression of *PECTIN METHYLESTERASE INHIBITOR 8* (*PMEI8*). These data suggest that *MYB50* is a new root growth regulator under the *UPB1* gene regulatory network, which differs from the control of ROS homeostasis. Our study presents a model including a new transcriptional network under MYB50 into UPB1 regulatory root growth system and will provide novel insights into the cell elongation controlled by pectin modification.

## Introduction

Plant roots are important organs that support the entire plant body, absorb water and nutrients from the soil, and respond to environmental changes. To achieve sufficient root architecture, the root developmental mechanism is strictly controlled. Maintaining the balance of transition from cell proliferation to differentiation at the root tip is the key to correcting primary root growth. Cell proliferation occurs in the meristematic zone that includes the stem cell niche. Once the cell leaves the meristematic zone, it stops proliferating but starts rapidly elongating in the elongation zone. After the cells fully elongate, they start to differentiate; in turn, the cells mature into their final states and function in the maturation zone. The transcriptional network is important for the transition to cell status. Several key transcription factors that regulate stem cell function have been identified in the meristematic zone. One important transcription factor is *WUSCHEL-LIKE HOMEOBOX5* (*WOX5*). *WOX5* acts as the key regulator for maintaining quiescent center (QC) cells [1, 2]. The second is *PLETHORA* (*PLT*). PLT proteins are master regulators of the meristematic zone [3–5]. PLT proteins are regulated by protein levels, and this regulation is important for determining cell function in the proliferative state [6]. These two key regulators interact with the auxin signaling pathway [7]. Auxin also creates a transcriptional circuit for defining meristem size with transcription factors, such as *ARR1* which is involved in cytokinin signaling [8]. Moreover, PLT proteins stability are regulated by the peptide hormone, ROOT GROWTH FACTOR (RGF) [9]. In addition to auxin and RGF, jasmonic acid regulates *PLT* through the transcription factor *MYC2* [10]. Ethylene regulates the QC status by regulating *ERF115* [11]. Braisnosteroids are directly regulated by cell cycle progression via *BES1* signaling [12]. Almost all plant hormones regulate the balance between cell proliferation and differentiation at the root tips. In addition to plant hormones, reactive oxygen species (ROS) also act as regulators to maintain the balance of cell function in the root tip; ROS, such as superoxide, hydrogen peroxide, and hydroxyl radicals, are thought of as molecules that are harmful to the cells since they have highly reactive potential. However, plant roots accumulate ROS at some levels to control root growth [13]. Hydrogen peroxide represses the expression of cell cycle-related genes in the meristematic zone [14]. *UP BEAT1* (*UPB1*), a bHLH domain-containing transcription factor, is a key regulator of the balance between cell proliferation and differentiation by regulating the expression of several peroxidase genes. The spatial accumulation patterns of superoxide and hydrogen peroxide control the size of the meristematic and elongation zones [15]. We have also reported that *MYB30* is a ROS-responsible transcription factor and *MYB30* regulates root cell elongation under ROS and ABA signaling [16, 17]. ANAC032 regulates *MYB30* expression and the transcription network that controls cell elongation [18]. These results indicate that ROS regulates both cell proliferation and elongation. Interestingly, *MYB30* and *ANAC032* are not the direct targets of UPB1. *ERF115*, *ERF114*, and *ERF109* respond to ROS and ethylene, and regulate cell proliferation and differentiation independent of PLT regulation [19]. Recently, a transcription factor, *RGF1 INDUCIBLE TRANSCRIPTION FACTOR 1* (*RITF1*), which controls ROS distribution at the root tip, was identified. The change in ROS distribution controlled by RITF1 enhances PLT2 protein stability and regulates meristematic zone size [9]. These results indicate that ROS signaling controls root development by integrating with other important regulatory networks for plant root development. Recent studies have revealed that ROS are important signaling molecules and indicate several targets of ROS signaling that regulate root growth. In other words, ROS signaling that regulates root growth has not yet been fully elucidated.

In the present study, we focused on the UPB1 gene regulatory network, which regulates primary root growth. Although UPB1 directly regulates the expression of several peroxidase genes, it still targets other genes. We previously identified 2,375 UPB1-responsive genes by microarray analysis and 166 putative UPB1 direct targets by ChIP-chip analysis. To elucidate UPB1 regulatory transcriptional network, we focused on the transcription factors among the 166 UPB1 direct targets. Among them, we chose the transcription factor, *MYB50* whose function has not yet been analyzed. We confirmed the transcriptional network from *UPB1* to *MYB50* by multi-color time-lapse imaging of *Arabidopsis* roots. We also investigated MYB50 regulatory networks using RNA-seq and ChIP-seq analyses. Based on these analyses, we found that MYB50 directly regulated the expression of several cell wall modification genes. We further investigated the root phenotypes of the MYB50 target genes, especially *PECTIN METHYLESTERASE INHIBITOR 8* (*PMEI8*). *PMEI8* overexpression resulted in a shorter mature cell length in the root. These results indicate that UPB1 regulates root growth through *MYB50* expression as a transcription factor for controlling further downstream gene expression and that MYB50 regulates cell elongation by controlling cell wall modification genes under the UPB1 transcription network.

## Material and Methods

### Plant materials and growth conditions

*Arabidopsis thaliana* Columbia-0 (Col-0) was used as the wild-type. The T-DNA insertion line, *upb1-1* was obtained from the SLAK collection and the seed stock center of the Arabidopsis Biological Resource Center (ABRC). *upb1-1* mutant was genotyped using a left-border primer on T-DNA (LB), right-side primers on the genome (RP), and left-border primers on the genome (LP) as described previously [15].

All seeds were sterilized with 1% bleach and 0.05% Triton X-100 for 5 min and then washed thrice with sterilized water. Seeds were germinated on Murashige and Skoog (MS; FUJIFILM Wako Pure Chemical, Osaka, Japan) medium supplemented with 1% sucrose and 1% agarose after two days at 4°C. Plants were grown vertically in a chamber (Panasonic, Osaka, Japan) at 22°C with 16-hour-light/8-hour-dark cycle. For estradiol treatment, 6-day-old seedlings were transferred onto MS agarose plates containing 5 µM estradiol (FUJIFILM Wako Pure Chemical).

### Plasmid construction and Plant transformation

Genomic DNA from Col-0 was used as a template for the amplification of the 3,020 bp upstream region of *pMYB50* for promoter cloning. One base of 5’ dA overhang was added to the PCR amplicon of *pMYB50* using Taq polymerase (Takara Bio Inc., Shiga, Japan), which was then cloned into pENTR5’-TOPO (Thermo Fischer Scientific, Waltham, MA, USA) and named pENTR5’-*pMYB50*. For the genomic coding region and cDNA for *MYB50* and *PMEI8* cloning, *MYB50* genomic and cDNA regions and the cDNA region of PMEI8 were amplified using the forward primer containing CACC sequences for TOPO cloning and the reverse primer. *CACC-gMYB50*, *CACC-cMYB50*, and *CACC-cPMEI8* fragments were cloned into pENTR/D-TOPO (Thermo Fischer Scientific). For *YFP-cMYB50*, *YFP-cPMEI8, YFP-cUPB1* and *CFP-cUPB1* cloning, the *cMYB50*, *cPMEI8*, and *cUPB1* cDNA regions were amplified using forward and reverse primers containing the BamHI site just before the *MYB50*, *PMEI8*, or *UPB1* termination codon. The *cMYB50*-Bam HI, *cPMEI8*- Bam HI, and *cUPB1*-Bam HI fragments were cloned into Aor51H1, and the Bam HI sites of *YFP*-Aor51H1- Bam HI-pDONR201 plasmids [16] and *CFP*-Aor51H1-Bam HI-pDONR201 plasmids [20], respectively.

For the *pMYB50::gMYB50-GFP* construct, pENTR5’-*pMYB50* and *gMYB50* containing pENTR/D-TOPO were cloned into R4pGWB650 [21] using LR Clonase II (Thermo Fischer Scientific). For the *pXVE::YFP- cMYB50*, *pXVE::YFP-cPMEI8*, *pXVE::YFP-cUPB1*, and *pXVE::CFP-cUPB1* constructs, *YFP-cMYB50*, *YFP- cPMEI8*, *YFP-cUPB1* and *CFP-cUPB1* containing pDONR201s were cloned into pMDC7 [22] using LR clonase II.

The resulting plasmids (pMYB50::gMYB50-GFP, pXVE::YFP-cMYB50, pXVE::YFP-cPMEI8, pXVE::YFP- cUPB1, and pXVE::CFP-cUPB1) were transformed into Agrobacterium tumefaciens (C58C1 pMP90) cells and transformed into Col-0 and upb1-1 mutants. For the pXVE::CFP-cUPB1/pMYB50::gMYB50-GFP double-reporter line, pXVE::CFP-cUPB1 containing Agrobacterium tumefaciens (C58C1 pMP90) cells were transformed into pMYB50::gMYB50-GFP plants. The sequences of primers used in this study are provided in S1 Table.

### Quantitative real-time RT-PCR

RNA was isolated from the whole roots of seven-day-old Col-0, and whole roots of six-day-old Col-0, *pXVE::YFP-UPB1*, *pXVE::YFP-MYB50*, and *pXVE::YFP-cPMEI8* plants treated with control MS and 5 µM estradiol for 24 h using the RNeasy Plant Kit (QIAGEN, Hilden, Germany). For RNA isolation from meristematic and elongation zones, six-day-old Col-0 and *upb1-1* mutants were microdissected as described previously [15]. First-strand cDNA was synthesized using ReverTra Ace qPCR RT Master Mix with gDNA Remover (TOYOBO Co., Ltd., Osaka, Japan). Quantitative real-time RT-PCR (RT-qPCR) was performed using THUNDERBIRD SYBR qPCR Mix (TOYOBO) on a real-time PCR Eco system (PCRmax, Stone Staffordshire, UK). The sequences of primers used in this study are provided in S1 Table. The RT-qPCR efficiency and CT values were determined using standard curves for each primer set. The efficiency-corrected transcript values of three biological replicates for all samples were used to determine the relative expression values. Each value was normalized against the level of *PDF2* [23].

### RNA-seq experiments

Total RNA was isolated from whole roots of Col-0 and *pXVE::YFP-MYB50* plants treated with 5 µM estradiol for 0, 1, 3, and 6 h. cDNA libraries were generated from 500 ng of total RNA using the NEBnext Ultra II RNA Library Prep kit (New England Biolabs), following the manufacturer’s protocols. The ends of the cDNA libraries were sequenced for 60 cycles using a paired-end module on the Illumina NextSeq 500 platform (Illumina, San Diego, CA, USA). Two biological replicates were used in each experiment.

### RNA-seq data analysis

Short-read sequencing results were mapped to the Arabidopsis genome (TAIR10: www.arabidopsis.org/) using the Bowtie software [24]. These datasets were normalized, and the False Discovery Rate (FDR) and Fold Change (FC) were calculated using the edgeR package for R [25] We used an FDR of *q* < 0.001 as the cut-off to determine differentially expressed genes between Col-0 and *pXVE::MYB50-GFP* treated with 5 µM estradiol. The data were deposited in the DNA Data Bank of Japan (DDBJ) sequence Read Archive (DRA) (https://www.ddbj.nig.ac.jp/index-e.html) under accession number DRA016078.

### ChIP-seq experiments

Over 1,200 *pXVE-YFP-MYB50* /Col-0 and Col-0 plants were grown on MS medium for 6 days. These plants were then transferred onto MS medium containing 5 µM estradiol. One day after transfer, whole roots were fixed and flash-frozen by liquid nitrogen according to a previously described protocol [15]. ChIP was performed as described previously [26] with anti-GFP antibody (ab290, Abcam, Cambridge, UK). ChIP DNA library was generated with a ChIP-Seq Sample Prep Kit (Illumina, San Diego, CA, USA) and the resulting DNA library was analyzed by GAII (Illumina) as described [26].

### ChIP-seq data analysis

Base-calling of sequence reads obtained by GAII was performed using the GAII pipeline software. Mapping of these sequence reads was performed using Bowtie [24] with default parameters. The resulting sequence alignment/map (SAM) file was converted into a binary alignment/ map (BAM) format file by Samtools 0.1.18 [27]. Significant ChIP DNA peaks (FDR *q* < 10^-20^) were annotated as YFP-MYB50 binding loci using the Model-based Analysis of ChIP-Seq (MACS2) software [28], with the genome size parameter dm (1.2e8). The forward- and reverse-peak distributions were validated using MACS2 and visualized using R (http://www.R-project.org/). BAM and indexed BAM files were used to visualize mapping patterns using Integrative Genomics Viewer 2.13.0. Based on the sequence data of 1,000 bp upstream from the translation start site for each gene, obtained from the TAIR database (TAIR10), genes near each peak were identified using a local BLAST tool [29] with the following parameters: blastn -evalue 0.1 -outfmt 6. Manual curation was performed to remove incorrect annotations arising from similarities in promoter sequence regions. The ChIP-seq data were deposited in the DNA Data Bank of Japan (DDBJ) sequence Read Archive (DRA) (https://www.ddbj.nig.ac.jp/index-e.html) under accession number DRA016088.

### Phenotypic and microscopy analysis

To measure the whole root length, the roots were scanned using a flatbed scanner GT-7400U (Epson, Nagano, Japan) while growing on plates. The root length was measured by importing the scanned images into Fiji (imagej.net/software/fiji/). Laser scanning confocal microscopy of *pMYB50::gMYB50-GFP*, *pXVE::CFP- cUPB1*/*pMYB50::gMYB50-GFP*, *pXVE::YFP-cMYB50*, and *pXVE::YFP-cPMEI8* was performed using a Leica SP8 system (Leica Camera AG) with propidium iodide (PI, FUJIFILM Wako Pure Chemical). Roots were stained with PI in a 10 µgmL-1 dilution in water for 3–5 minutes, with 448 nm excitation and 460–510 nm emission for CFP, 488 nm excitation and 500–550 nm emission for GFP, 488 nm excitation and 490–543 nm emission for YFP, and 555 nm excitation and 580–680 nm emission for PI. Image assembly was performed using the LAS X software (Leica Camera AG).

For time-lapse imaging, Lab-Tek Chambered Coverglass w/cvr (Thermo Fisher Scientific) was used, as described previously [16]. For time-lapse imaging of multi-color reporter line, 6-day-old seedlings of *pXVE::CFP-cUPB1*/*pMYB50::gMYB50-GFP/upb1-1* in MS medium were placed on chambered cover glass in media containing 5 µM estradiol. The chambers containing plants were imaged using a Leica SP8 confocal microscope at objectives of 20x. Time-lapse images were captured using LAS X every 40 min for 7 h and 20 min. Images were also assembled with LAS X. To quantify CFP-cUPB1 and gMYB50-GFP intensities, 50 z-stack images of the roots were taken with LAS X every 20 min for 5 h using 40x (oil) objectives. Of the 50 z-stack images for each time point, the image with the most focus on the nucleus of the cortex or epidermal cells was selected, and CFP and GFP intensities were quantified as a mean intensity using Fiji software.

To measure the increase in root length, 4-day-old seedlings of Col-0, *pXVE::YFP-UPB1,* and *pXVE::YFP- MYB50* in MS medium were placed on chambered cover glass in media containing 5 μM estradiol. The chambers containing the plants were imaged using a DMI 6000 B-AFC fluorescence microscope (Leica Camera AG, Wetzlar, Germany) under a 20x objective lens. Time-lapse images were captured every 30 min for 24 h. The increase in root length was measured using the Fiji software on a time series consisting of images taken every 30 min.

### Statistical analysis

All statistical analyses were performed using Microsoft Excel or R. Hypergeometric testing for overlapping genes between ChIP and RNA-seq was performed using the online program http://nemates.org/MA/progs/overlap_stats.html. Details of the analyses are provided in the figure legends.

## Results

### UPB1 regulates MYB50 expression at the elongation zone

To confirm our previous results, we analyzed *MYB50* expression in the *upb1-1* mutant by RT-qPCR using RNA extracted independently from meristematic and elongation zones. Microdissection was confirmed to have been performed correctly by checking the expression of *CYCB1;1* and *UPB1*, which are known to be expressed in the meristematic and elongation zones, respectively (Fig 1A). *MYB50* expression was significantly higher in the elongation zone of the *upb1-1* mutant than in Col-0 plants (Fig 1A). Consistent with our previous results, UPB1 suppressed *MYB50* expression in the elongation zone [15]. We constructed an estradiol-inducible *YFP-UPB1* transformant (*pXVE::YFP -UPB1*). After 24 h of 5 µM estradiol treatment, *UPB1* expression was strongly induced and *MYB50* decreased significantly (Fig 1B). Time-lapse imaging was performed every 30 min after estradiol treatment to measure root elongation. *pXVE::YFP-UPB1* showed that root elongation was strongly inhibited after estradiol treatment compared to Col-0 (Fig 1C). These results indicated that *UPB1* induced-overexpressor repressed *MYB50* and affected root growth, similar to our previous results [15].

**Fig 1.**
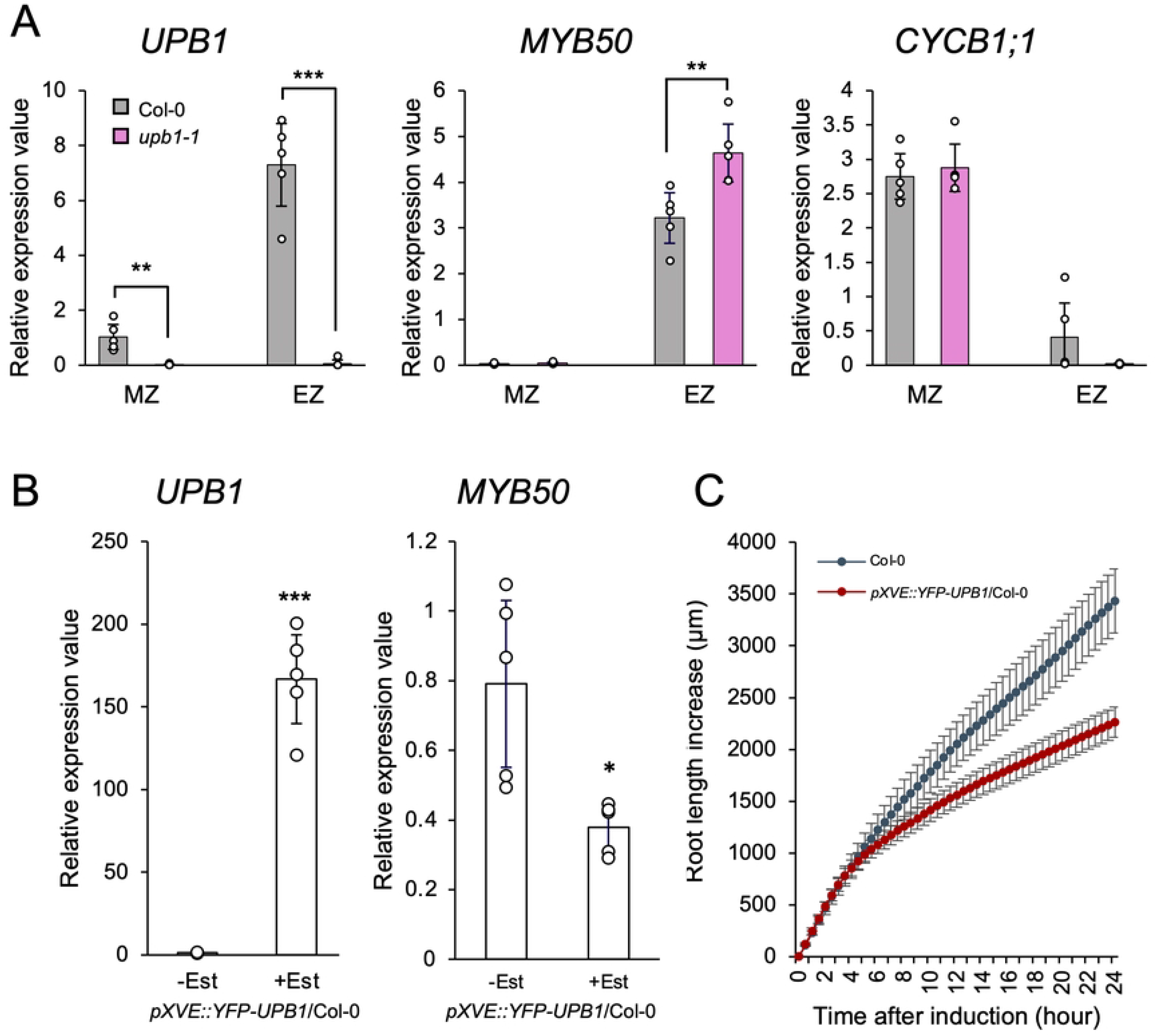
UPB1 regulates *MYB50* expression at the elongation zone. (A) *UPB1*, *MYB50,* and *CYCB1;1* expression in both of the meristematic and elongation zones in Col-0 (gray) and *upb1-1* (pink) as measured by RT-qPCR (n = 5). MZ and EZ indicate the meristematic and elongation zone, respectively. Bars show mean ± SD. Significant differences were determined by Student’s *t*-test by comparing Col-0 and *upb1-1* in each zone (*** *p* < 0.001; ** *p* < 0.01; * *p* < 0.05). *p* value: comparison of *UPB1* expression in MZ of Col-0 and *upb1- 1*, *p* = 0.002; comparison of *UPB1* expression in EZ of Col-0 and *upb1-1*, *p* < 0.0001; comparison of *MYB50* expression in EZ of Col-0 and *upb1-1*, *p* = 0.0096. (B) UPB1 and MYB50 expression analysis of whole roots in *pXVE::YFP-UPB1*/Col-0 with and without 5 μM estradiol treatment for 24 h measured by RT-qPCR (n = 5). Significant differences were determined by Student’s *t*-test compared with and without estradiol treatment (*** *p* < 0.001; * *p* < 0.05). *p* value: comparison of *UPB1* expression, *p* < 0.001; comparison of *MYB50* expression, *p* = 0.016. (C) Root length increase in 4-day-old seedlings of Col-0 and *pXVE*::*YFP-UPB1*/Col-0 treated with 5 μM estradiol containing agarose medium. Root length increase for 24 hours was measured every 30 minutes by time-lapse imaging (n = 5, means ± SD).

To investigate *MYB50* expression patterns in roots, we constructed a translational fusion reporter of *MYB50* (*pMYB50::gMYB50-GFP*) in Col-0 or *upb1-1* mutant backgrounds. In *MYB50* translational fusion reporter in the Col-0 background, MYB50-GFP signals were detected in the early differentiation zone but not in the meristematic and elongation zones. Moreover, *MYB50* was expressed in the epidermis, cortex, endodermis, and pericycle cells (Fig 2A). However, *MYB50* translational fusion reporter in the *upb1-1* mutant showed a GFP signal in the nucleus of the elongation zone but not in the meristematic zone (Fig 2A). This result also indicated that *MYB50* was regulated by UPB1 in the elongation zone. Then, we performed time- lapse imaging by using two fluorescent proteins. For time-lapse imaging, we fused estradiol-inducible *UPB1* with *CFP* (*pXVE::CFP-UPB1*) and transformed *pXVE::CFP-UPB1* into the *pMYB50::gMYB50-GFP/upb1-1* reporter line. Because GFP and CFP possess different emission wavelengths, we could detect the fluorescence from GFP and CFP simultaneously. This allowed us to detect the spatiotemporal transcriptional networks in the same cell as the root (S1 Movie). Transcription factors like UPB1 and MYB50 are localized in the nucleus. For quantification of the fluorescence intensity, the problem of signal detection was focused on during time-lapse imaging because of nuclear movement in the cells. Therefore, we performed z-stack time-lapse imaging, focusing on cells that expressed MYB50 in the early differentiation zone. Among the 50 z-stack images, the image with the greatest focus on the nucleus was selected for each time point, and both CFP-cUPB1 and gMYB50-GFP intensities were quantified (Figs 2B, 2C, and S1 Fig). After 180 minutes of 5 µM estradiol treatment, CFP-UPB1 started to appear strong and GFP fluorescence was almost unaltered (Fig 2B and 2C). After 220 min of estradiol treatment, GFP fluorescence was weaker than that at the initial time points (Fig 2B and 2C). Fluorescence intensity was also quantified. As CFP intensity increased, GFP expression weakened (Fig 2C). Moreover, fluorescence alterations occurred between the elongation and earlier differentiation zones. Supporting this result, the mRNA levels of *UPB1* and *MYB50* in this multi-color reporter line were significantly altered after 5 h of estradiol treatment, similar to the fluorescence intensity (S2 Fig). These results strongly indicate that MYB50 is one of the UPB1 direct target transcription factors that mediated UPB1 signaling for root growth and that UPB1 represses *MYB50* expression in the elongation zone.

**Fig 2.**
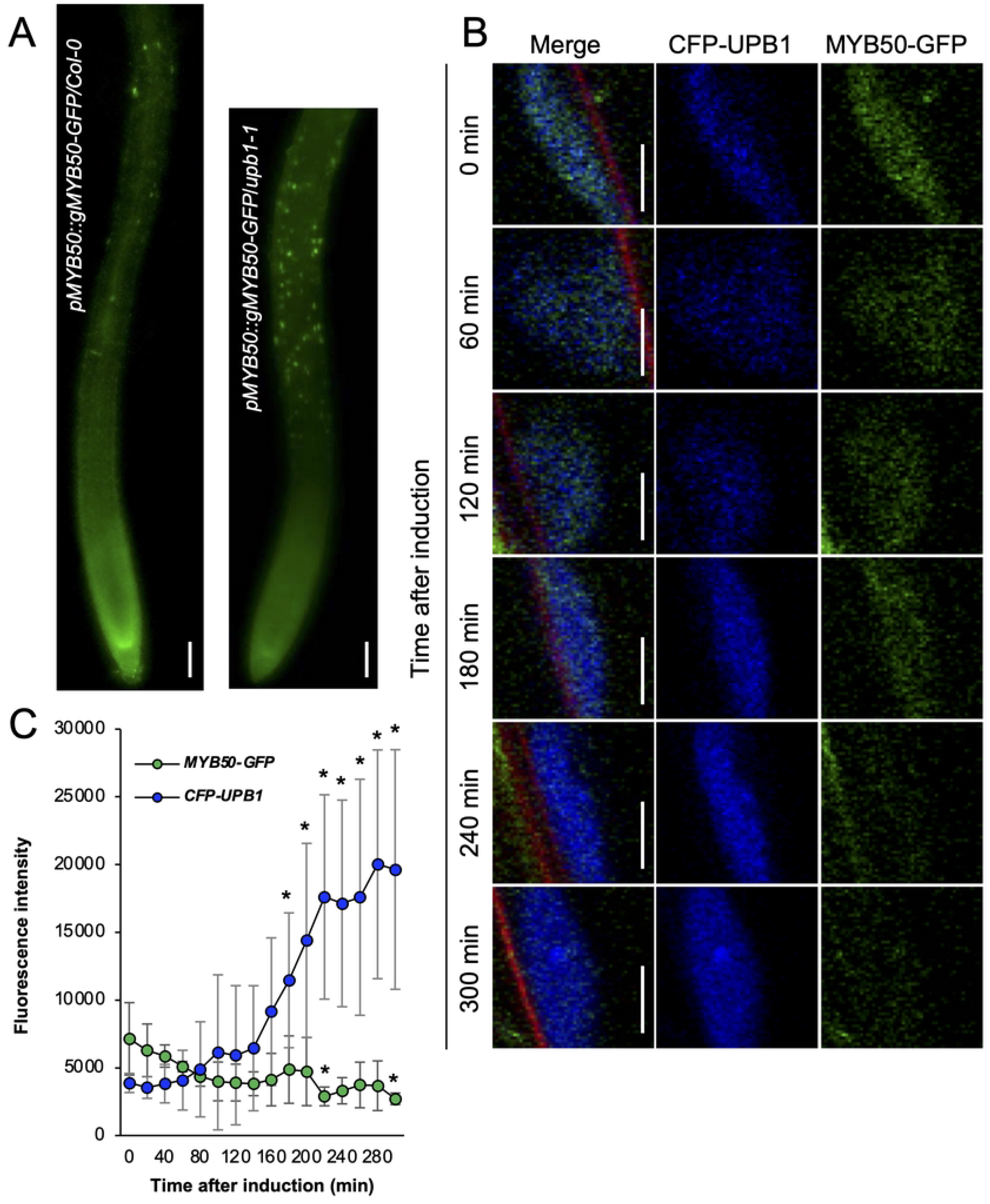
*MYB50* expression pattern in the primary root and the multi-color fluorescent protein expression of CFP-UPB1 and MYB50-GFP. (A) MYB50-GFP expression in the primary root tip of *pMYB50::gMYB50- GFP*/Col-0 and *pMYB50::gMYB50-GFP*/*upb1-1*. Scale bars, 100 µm. (B) Timepoint images every 60 minutes in the visualization of transcriptional repression of MYB50 by UPB1 in one nucleus. 6-day-old seedlings of *pXVE::CFP-UPB1*/*pMYB50::gMYB50-GFP*/*upb1-1* were stained by propidium iodide (PI) and CFP-UPB1 and each MYB50-GFP signal was independently detected. Left panels: merged images of CFP, GFP, and PI, Middle panels: CFP channel, Right panels: GFP channels. Scale bars, 5 µm. (C) Quantification of CFP-UPB1 and MYB50-GFP intensity after induction every 20 minutes. Each signal intensity was measured every 20 minutes by time-lapse imaging (n = 4, means ±SD). Significant differences for GFP and CFP were determined by Student’s *t*-test compared to 0 minute-induction (* *p* < 0.05). *p* value: GFP after 220 minutes, *p* = 0.0374; GFP after 300 minutes, *p* = 0.0299; CFP after 200 minutes, *p* = 0.04; CFP after 220 minutes, *p* = 0.0447; CFP after 240 minutes, *p* = 0.0241, CFP after 260 minutes, *p* = 0.0348; CFP after 280 minutes, *p* = 0.0164, CFP after 300 minutes, *p* = 0.0218.

### MYB50 regulates the size of the meristematic zone and mature cell length

After showing that *MYB50* is in the *UPB1* transcriptional network, we next investigated how *MYB50* is involved in root growth. We attempted to analyze the *MYB50* function in root growth using genetic methods but the *MYB50* T-DNA insertion line did not exist in the ABRC seed stock center. To investigate the effects of *MYB50* overexpression on root growth, we made an *MYB50* fused with a YFP estradiol-inducible line (*pXVE::YFP-MYB50*). In this inducible line, *MYB50* expression reached more than 100 times the Col-0 level in both independent lines after 24 h of estradiol treatment (Fig 3A). We investigated the root growth phenotype caused by induced *MYB50* with time-lapse imaging every 30 min for 24 h. After 24 h of induction with 5 μM estradiol, root elongation of *pXVE::YFP-MYB50* was inhibited (Fig 3B). To determine the detailed effects of *MYB50* on root growth, we counted the number of cortical cells in the meristematic zone and measured the mature cell length in *MYB50* induced-overexpressor. Two independent lines of *pXVE::YFP- MYB50* treated with 5 μM estradiol for 24 h showed reduced cortical cell numbers in the meristematic zone and shorter mature cell length (Fig 3C-3F). Taken together, these results suggest that *MYB50* overexpression negatively regulates meristematic zone size and mature cell length, similar to *UPB1* overexpression. Because *UPB1* is known to control ROS homeostasis at the root tip, we investigated how the inhibition of root elongation under MYB50 regulation and ROS-controlled root growth interacted with each other. H_2_O_2_, a ROS, is known to inhibit root elongation [30]. The combination treatment with estradiol and H_2_O_2_ resulted in a smaller meristem size than estradiol treatment alone in Col-0 cells (S3 Fig). Interestingly, *MYB50* induced- overexpressor showed smaller meristem sizes than Col-0 plants in the estradiol treatment but even smaller meristem sizes upon H_2_O_2_ and estradiol treatment (S3 Fig). This additive effect suggests that the MYB50 regulatory network inhibits root growth independent of ROS regulation.

**Fig 3.**
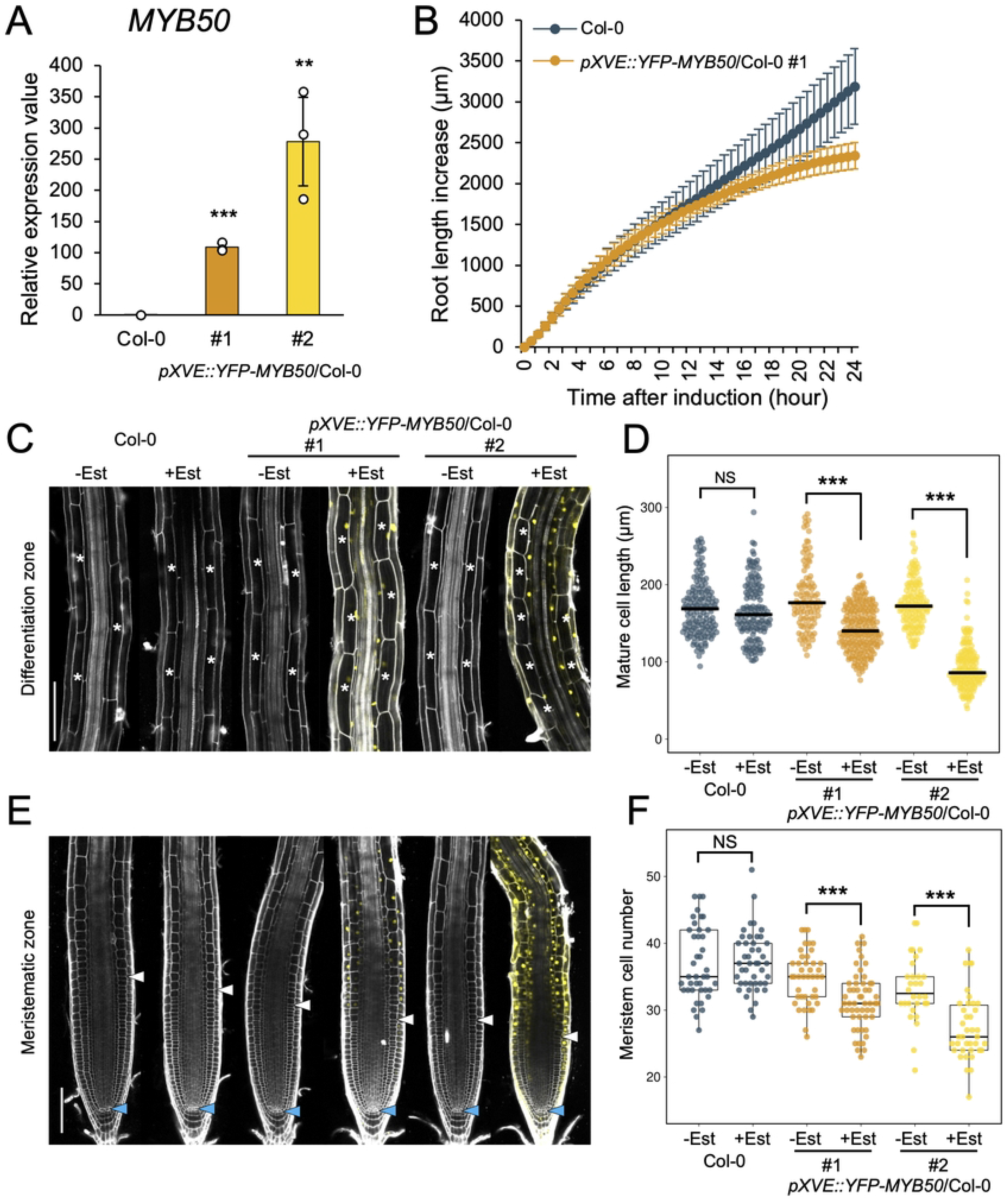
Root phenotype of *MYB50* induced-overexpressor (*pXVE::YFP-MYB50*/Col-0). (A) *MYB50* expression in Col-0 and two independent lines of *pXVE::YFP-MYB50*/Col-0 after 24 h treatment of 5 μM estradiol (Est) as measured by RT-qPCR (n = 3, mean ± SD). Significant differences were determined by Student’s *t*-test compared to Col-0 (*** *p* < 0.001; ** *p* < 0.01). *p*-value: comparison of Col-0 and #1, *p* < 0.001; comparison of Col-0 and #2, *p* = 0.0052. (B) Root length increase in 4-day-old seedlings of Col-0 and *pXVE::YFP-MYB50*/Col-0 treated with 5 µM Est containing agarose medium. Root length increase was measured every 30 minutes by time-lapse imaging (n = 5, means ± SD). (C) and (E) Confocal microscope images of Col-0 and *pXVE::YFP-MYB50*/Col-0 that were treated with 5 μM Est for 24 h or were untreated. Roots were stained with propidium iodide. (C) The differentiation zone is at a distance of about 1 to 2 mm from each root tip. White asterisks indicate cortex cells with cell length measured. Scale bar, 100 µm. (D) Length of mature cortex cells in the Col-0 and *pXVE::YFP-MYB50*/Col-0 (two independent lines). The black bar indicates the median. Mature cell length was quantified for 10 to 15 roots, with approximately 3 to 10 cells per root. Measured cell numbers: Col-0 -Est = 149, Col-0 +Est = 141, *MYB50* induced-overexpressor #1 -Est = 101, #1 +Est = 209, #2 -Est = 130 and #2 +Est = 150. Significant differences were determined by Student’s *t*-test compared Est treated and untreated (*** *p*< 0.001). (E) The meristematic zone in Col-0 and *MYB50* induced-overexpressor with 5 μM Est treatment for 24 hours or no treatment. White arrowheads indicate the end of the meristematic zone. Blue arrowheads indicate quiescent cells (QC). Scale bar, 100 µm. (F) Cortex cell number in the meristematic zone treated with 5 μM Est or untreated Col-0 and *pXVE::YFP-MYB50*/Col-0. Counted root numbers: Col-0 -Est = 41, Col-0 +Est = 44, MYB50 induced-overexpressor #1 - Est = 42, E#1 +Est = 53, #2 -Est = 30, #2 +Est = 38. Significant differences were determined by Student’s *t*- test compared to Est treated and untreated plants (*** *p* < 0.001).

### Identification of MYB50-regulated genes

To identify the MYB50 gene regulatory network, we conducted RNA-Seq and ChIP-Seq analyses. We assumed that analyzing estradiol-inducible transcription factors would be reliable because it would be possible to track the prompt effects of the overexpression of the target transcription factors controlled by estradiol treatment. In other words, rapid regulation triggered by the overexpression of inducible genes can identify the primary target of the transcription factor of interest without secondary or tertiary effects. We then performed time-course RNA-seq analysis of *pXVE::YFP-MYB50* roots treated with 5 μM estradiol for 1, 3, and 6 h. Based on these analyses, we found that 210, 90, and 141 genes (singletons) were significantly upregulated at 1, 3, and 6 h, respectively (two-fold change, FDR < 0.001; S2 Table). In contrast, 49, 54, and 96 genes (singletons) were downregulated at 1, 3, and 6 h, respectively (two-fold change, FDR < 0.001; S3 Table). We analyzed the Gene Ontology (GO) terms of the “Biological Process” of upregulated genes in *MYB50*-induction lines and found that 15, 19, and 11 GO terms were significantly enriched at 1, 3, and 6 h, respectively (FDR < 0.001; Fig 4A). Among enriched GO terms, “Response to acid chemical,” “Response to abiotic stimulus,” “Response to chemical,” and “Response to oxygen-containing compounds,” were highly enriched. This indicates that *MYB50* may be involved in stress responses.

**Fig 4.**
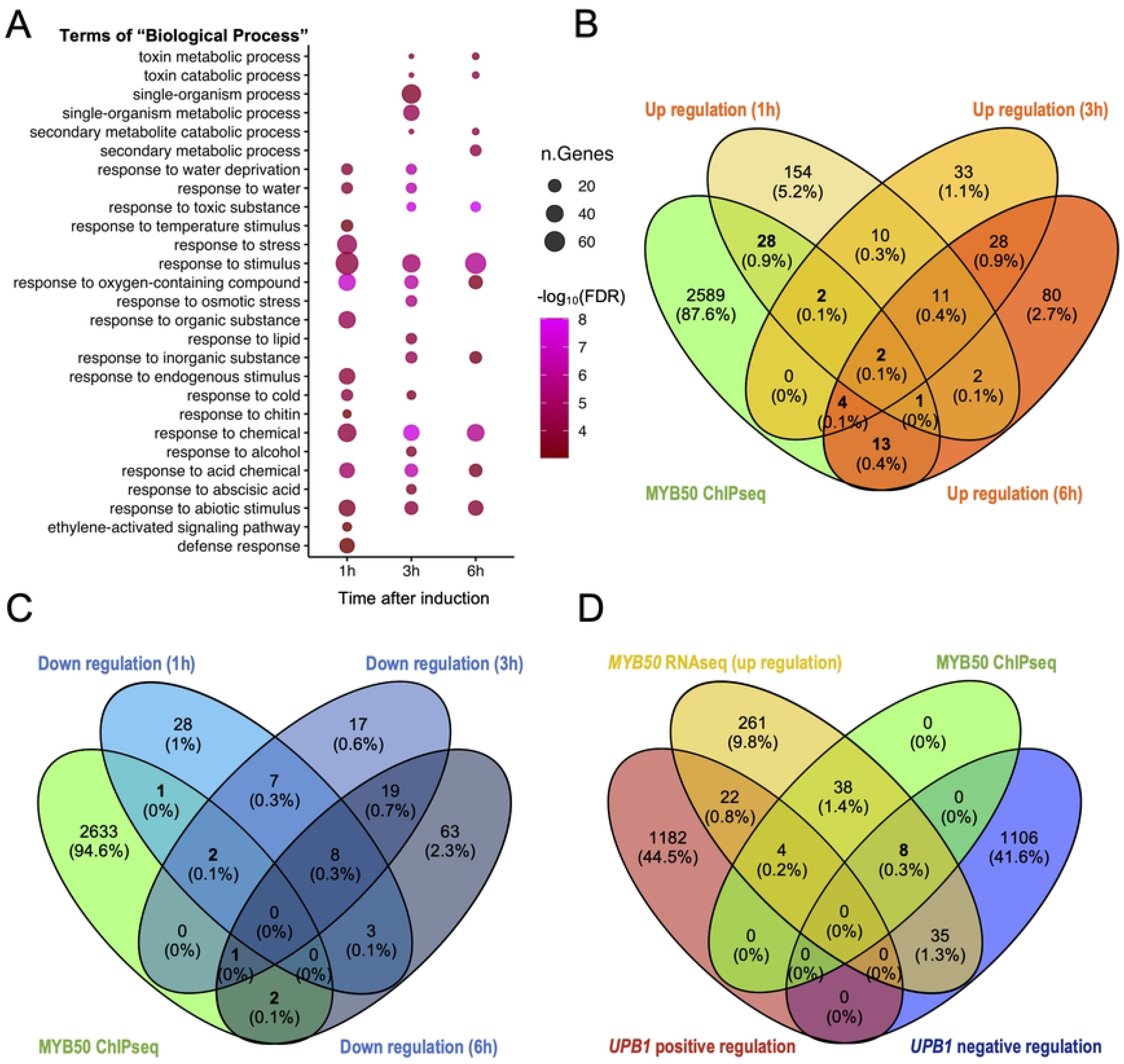
MYB50 regulated genes. (A) Biological processes in Gene Ontology (GO) enrichment analysis of upregulated genes in *MYB50* induced-overexpressor (FDR < 0.001). The color and size of each point represented the -log_10_ (FDR) values and the number of genes, respectively. (B) Overlapping genes between upregulated genes in each timepoint of RNA-Seq and MYB50-bound genes by ChIP-Seq. (C) Overlapping genes between downregulated genes in each timepoint of RNA-Seq and MYB50-bound genes by ChIP-Seq. (D) Overlapping genes between MYB50- and UPB1-regulated genes. *MYB50* RNA-Seq (upregulation): Upregulated genes (368 genes) in at least one-time point of estradiol induction of *pXVE::YFP-MYB50*/Col-0 in RNAseq. MYB50 ChIP-Seq: Overlapping genes (50 genes) between upregulated genes by MYB50 and MYB50-bound genes in *pXVE::YFP-MYB50*/Col-0. UPB1 positive regulation: Positively regulated genes (1,208) by UPB1. UPB1 negative regulation: Negatively regulated genes (1,149) by UPB1. UPB1 data were retrieved from Tsukagoshi et al., 2010 [15].

To identify MYB50 direct targets, we performed ChIP-seq analysis using 24 h estradiol-treated *pXVE::YFP-MYB50*. Sequence reads were mapped to the *Arabidopsis* genome TAIR10 (https://www.arabidopsis.org/index.jsp) and genomic regions bound by *MYB50* according to an FDR *q*-value of < 10^-20^ in the dataset (supplemental dataset 1). ChIP-seq analysis identified 2,639 regions 1,000 bp upstream of the translation initiation site of genes. Of course, this number included false-positive genes because of the technical limitations of the ChIP-Seq protocol. To find *MYB50*-biological direct target genes, we combined RNA-Seq and ChIP-Seq data. We searched for overlapping genes between MYB50-bound genes in ChIP-Seq and total genes that were upregulated (368 genes) or downregulated (151 genes) at least once during estradiol induction of *MYB50* in RNA-Seq. Overlapping genes (50 genes) between the upregulated genes and MYB50 bound genes were statistically significant (hypergeometric probability test, *P* = 3.23e-07), whereas overlapping genes (six genes) between the downregulated genes and MYB50 bound genes were not (hypergeometric probability test, *P* = 0.151; Fig 4B and 4C). This suggests that MYB50 preferentially upregulates genes bound to MYB50 in the promoter region.

Since *MYB50* was selected among UPB1 direct target genes, we compared differentially expressed genes (DEGs) between *UPB1* and *MYB50-regulated* genes (Fig 4D). Among the 368 genes upregulated in *pXVE::YFP-MYB50*, 43 were negatively regulated by UPB1 (1,149 genes). Eight genes were MYB50 ChIP-positive. In contrast, 26 genes were positively regulated by UPB1 (1,208 genes). Four genes were positive for MYB50 by ChIP (Fig 4D and S4 Table). Given that the number of overlapping MYB50- and UPB1-regulated genes was not very high may be due to transcriptional feedback regulation by the corresponding transcription factors. The eight genes that were direct targets of MYB50 and were negatively regulated by UPB1 included stress-and ABA-responsible genes, such as *Dehydrin family protein* (*ERD10*; *At1g20450*), *CBL-interacting protein kinase 9* (*CIPK9*; *At1g01140*), and *cold-regulated 47* (*COR47*; *At1g20440*), and cell wall modification genes, such as *Plant invertase/pectin methylesterase inhibitor superfamily protein* (*PMEI8*; *At3g17130*). Among these eight genes, *PMEI8* was the most strongly upregulated in the RNA-Seq data sets (S4 Fig).

### MYB50 regulates the expression of pectin modification genes

Among the *MYB50* direct target genes which also showed differentially expressed in *upb1-1*, we focused on the role of *PMEI8* (*At3g17130*) in root growth. Pectin methylesterase inhibitors (PMEIs) are known to regulate pectin status, and *PMEI8* overexpression is partially complemented by defective root growth in *cobra* mutants with a short-root phenotype [31]. However, the detailed effects of *PMEI8* overexpression on root growth have not been reported. In silico analysis of RootMap gene expression data [32] showed that *PMEI8* expression peaked in section 8 (the elongation zone close to the differentiation zone) in each tissue (S5 Fig). This expression pattern indicates that MYB50 regulates *PMEI8* expression in the elongation zone (Fig 2A). In addition, MYB50 bound approximately 1,000 bp upstream of the *PMEI8* coding region according to our ChIP-Seq data (Fig 5A). Since *PMEI8* was upregulated in the *MYB50* induced- overexpressor (Fig 5B), we decided to investigate the phenotype of the overexpressor of *PMEI8*.

**Fig 5.**
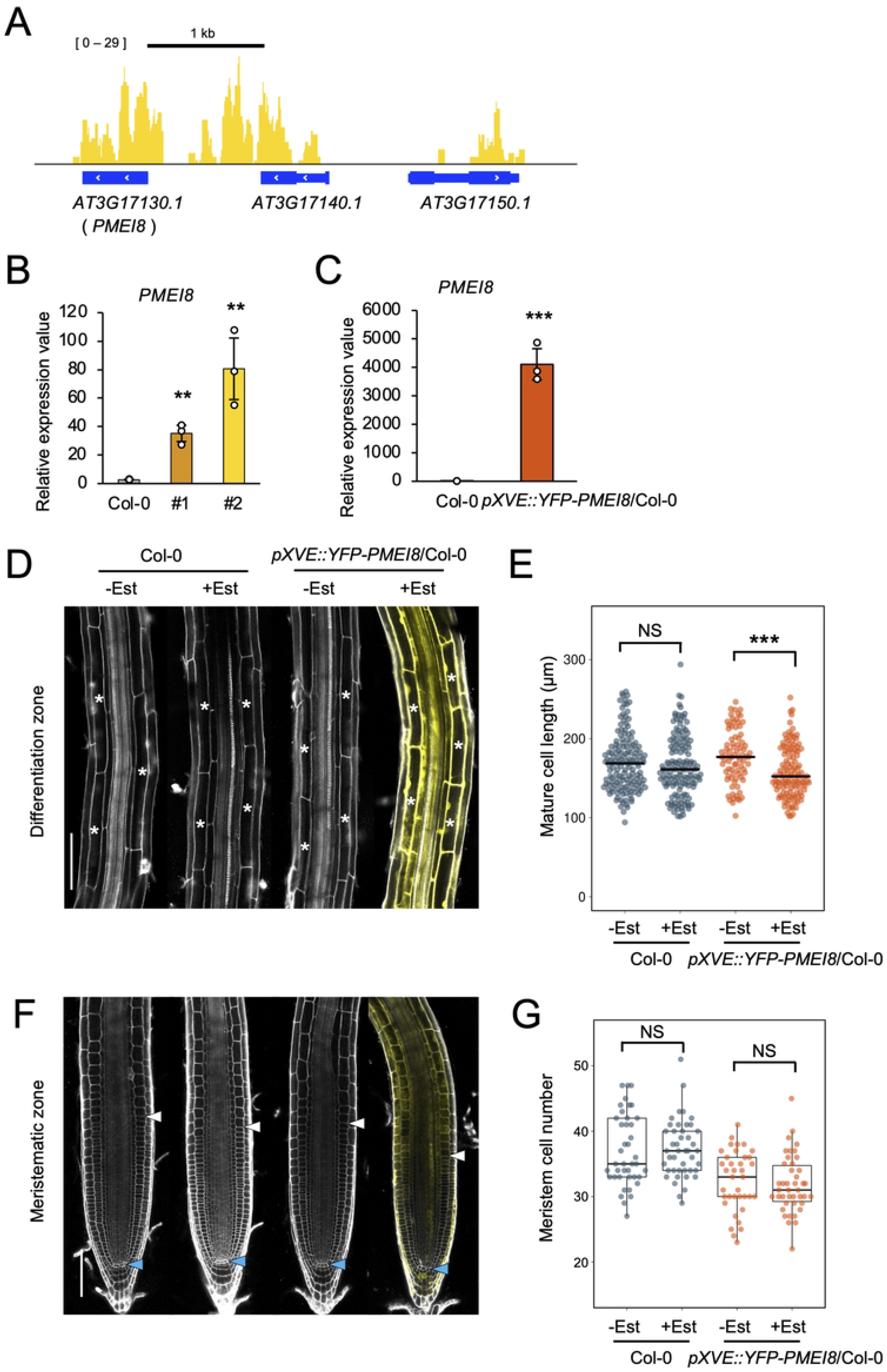
Root phenotype of *PMEI8* induced-overexpressor (*pXVE::YFP-PMEI8*/Col-0). (A) MYB50 bound to the *PMEI8* promoter region in ChIP-seq. “Blue boxes” and “white >” indicate ORF (thick box: exon; thin box: intron) and sequence toward (>: forward; <: reverse) respectively. (B) *PMEI8* expression of Col-0 and *pXVE::YFP-MYB50*/Col-0 (two independent lines) after 24 h of 5 μM estradiol (Est) treatment as measured by RT-qPCR (n = 3, mean ± SD). Significant differences were determined by Student’s *t*-test compared to Col-0 (** *p* < 0.01). *p* value: comparison of Col-0 and #1, *p* = 0.00131; comparison of Col-0 and #2, *p* = 0.00702. (C) *PMEI8* expression of *pXVE::YFP-PMEI8*/Col-0 after 24 h of 5 μM Est treatment as measured by RT-qPCR (n = 3, mean ± SD). A significant difference was determined by Student’s *t*-test compared to Col-0 (*** *p* < 0.001). (D) and (F) Confocal microscope images of Col-0 and *pXVE::YFP-PMEI8*/Col-0 that were treated with 5 μM Est for 24 h or untreated. Roots were stained with propidium iodide (PI). (D) The differentiation zone is at a distance of about 1 to 2 mm from each root tip. White asterisks indicate cortex cells with cell length measured. Scale bar, 100 µm. (E) Length of mature cortex cells in the Col-0 and *pXVE::YFP- PMEI8*/Col-0. Mature cell length was quantified for 10–15 roots, with approximately 3–10 cells per root. The black bar indicates the median. Measured cell numbers: Col-0 -Est = 149, Col-0 +Est = 141, *PMEI8* induced- overexpressor-Est = 81, *PMEI8* induced-overexpressor +Est = 120. Significant differences were determined by Student’s *t*-test compared to Est treated and untreated (*** *p* < 0.001). (F) The meristematic zone in Col-0 and *PMEI8* induced-overexpressor with 5 μM Est treatment for 24 hours or untreated. White arrowheads indicate the end of the meristematic zone. Blue arrowheads indicate the quiescent cells (QC). Scale bar, 100 µm. (G) Cortex cell number in the meristematic zone treated with 5 μM Est or untreated of Col-0 and *pXVE::YFP-PMEI8*/Col-0. Counted root numbers: Col-0 -Est = 41, Col-0 +Est = 44, *PMEI8* induced- overexpressor -Est=37, *PMEI8* induced-overexpressor +Est=42. Significant differences were determined by Student’s *t*-test compared to estradiol-treated and untreated plants.

To investigate how *PMEI8* overexpression affected root growth, we made a *YFP-PMEI8* inducible line (*pXVE::YFP-PMEI8*) that expressed *PMEI8* more than 4,000 times after 24 h of estradiol treatment (Fig 5C). We measured the number of cortical cells in the meristematic zone and the mature cell length in the *PMEI8* induced-overexpressor. The cell numbers in the meristematic zone of estradiol-treated *pXVE::YFP- PMEI8* were slightly but not significantly, reduced compared to those of Col-0. Estradiol-treated *pXVE::YFP- PMEI8* showed significantly shorter mature cell length than control cells (Fig 5D-5G). The mature cell length phenotype was consistent with *MYB50* induced-overexpressor in which MYB50 directly upregulated *PMEI8* expression (Fig 5A and 5B). This result indicates that *PMEI8* is a negative regulator of root growth, especially cell length, in the *MYB50* transcriptional network.

## Discussion

### *MYB50* is a UPB1 direct downstream transcription factor gene

In a previous study, we showed that UPB1 regulates root development through ROS homeostasis at the root tip by directly controlling the expression of peroxidase genes [15]. In the present study, we found that UPB1 binds to the promoter regions of several transcription factor genes that are not involved in ROS homeostasis. We hypothesized that UPB1 target transcription factors may also regulate root growth through a different regulatory network independent of ROS. Among these, we were interested in a novel transcription factor that has not yet been reported. In this study, we selected *MYB50* as a candidate gene that mediates the UPB1 regulatory network for root growth. *MYB50* expression was upregulated in the *upb1-1* mutant but downregulated in the UPB1 overexpressor. Since *MYB50* is expressed in the elongation zone, *MYB50* would have function in the root development. In support of this, *MYB50* induced-overexpressor showed decreased root elongation compared to the control. These results indicate that *MYB50* is a root growth inhibitor. We confirmed *UPB1*-*MYB50* transcriptional regulation by multi-color reporter lines. Visualizing transcriptional regulation is challenging; however, recent imaging technologies have allowed us to achieve this goal. MYB50-GFP expression was repressed after the estradiol induction of CFP-UPB1 in root cells. This system might be used not only for other gene regulatory networks of transcription factors but also for tracing alterations in the subcellular localization of several proteins simultaneously.

### MYB50 regulated *PMEI8* expression under UPB1 gene regulatory network

Transcriptome analysis of *MYB50* induction indicated that MYB50 regulates the expression of ROS homeostasis genes regulated by UPB1. In support of this, H_2_O_2_ treatment had an additive effect on the reduction in meristem size in *MYB50* inducible overexpressor. Interestingly, the number of overlapping MYB50 and UPB1-regulated genes was low. Since the *MYB50* expression level by the *pXVE* promoter was much stronger than that in *upb1-1* mutants, *MYB50* overexpression affected its downstream region more than the UPB1 gene regulatory network. Another possibility is the feedback regulation of *MYB50* overexpression or the *upb1-1* mutation. However, we found one interesting gene, *PMEI8*, among *MYB50* target genes which was also de-repressed in the *upb1-1* mutant. Our results showed that *PMEI8* under the MYB50 gene regulatory network inhibited cell elongation and consequently shortened mature cell length. PMEIs, including methyl esterified pectin, function as PME inhibitors. Methyl-esterified pectin stiffens pectin gels and inhibits cell elongation [33, 34]. Fine-tuning of pectin methyl esterification is thought to be key to cell wall stiffening, which regulates cell growth. Spatiotemporal control of pectin methyl esterification by PMEs and PMEIs in *Arabidopsis thaliana*. Although PMEI family proteins are predicted to have the same function, each PMEI can exhibit different root phenotypes. *PMEI9* overexpression resulted in a longer primary root phenotype, whereas *PMEI4* overexpression resulted in a short root phenotype [35]. Other *PMEI*s, such as *PMEI-1* and *PME-2*, have longer roots than the wild type [36]. These results about PMEI function support the hypothesis that spatiotemporally controlled PMEIs regulate intrinsic cell growth. Moreover, a single overexpression of another *PMEI* (*At5g62360*) did not alter root length; however, overexpression of *PMEI* with *EXPA5*, which is thought to be a cell wall-loosening enzyme, increased root length [37]. *PMEI8* has been identified as a partial suppressor of *cobra* mutations. *PMEI8* overexpression in *cobra* mutants can recover their root growth defects [31]. According to these functions, there are co-regulatory genes of PMEI8 in the *MYB50* and *UPB1* regulatory gene networks. To support this hypothesis, we identified several cell wall modification genes in the transcriptome datasets. To fully understand the *MYB50* gene regulatory network that controls root growth, we need to investigate the gene functions in the near future.

We found that 368 genes were significantly upregulated in the RNA-Seq analysis of the *MYB50* induction line. Among the upregulated genes, we identified GO terms representing the responses to abiotic stress and chemical compounds (Fig 4A). Furthermore, we combined these data with ChIP-Seq data from YFP-MYB50, concluded that *MYB50* was a transcriptional activator, and identified 50 genes as MYB50 direct target genes. We identified *ABA repressor 1* (*ABR1*), *NAC004*, *ERF019*, and *CBF2* [38–39]. These transcription factors are all involved in the ABA response, especially in stress responses such as cold and drought. In addition to these transcription factors, we identified ABA and stress response marker genes such as *LEA*, *RD29A*, *ERD4*, *ERD10*, and *COR47* [40–46]. These results indicated that MYB50 may be involved in plant stress responses regulated by ABA. In future studies, to deepen our understanding of the function of MYB50 in ABA signaling, we need to investigate the phenotypes of *MYB50* induced-overexpressor under abiotic stress.

## Conclusions

In this study, we identified a transcription factor, *MYB50*, that acts as a mediator of the UPB1 gene regulatory network for regulating root growth. Gene expression analysis and multi-color imaging strongly indicate that *MYB50* expression is regulated by UPB1. The timing of switching off *MYB50* expression by UPB1 was coordinated with alterations in root growth. Based on RNA-seq and ChIP-seq analyses, we identified several co-regulated genes in the UPB1 and MYB50 gene regulatory networks. We found that *PMEI8* controlled root growth. The data presented above, combined with previous UPB1 data, are shown as a model of the gene regulatory network (Fig 6). UPB1 negatively regulates meristem size via ROS homeostasis, by repressing the expression of several peroxidase genes. *Peroxidase* overexpressors have large meristem sizes [15]. In this study, we showed that UPB1 directly represses *MYB50* in the elongation zone. MYB50 negatively regulated meristem size in the same direction as UPB1 since *MYB50-induced* overexpressor showed smaller meristem size and mature cell length (Fig 3E and F). It is still unclear which mediator suppresses meristem size under *MYB50* transcriptional regulation, although it has been revealed that *UPB1* gene regulatory network has two different regulatory genes, the accelerator and decelerator of meristem size. In the base from the meristematic zone, UPB1 suppresses cell elongation to a longitudinal direction [15]. We showed that MYB50 also suppressed cell elongation through transcriptional activation of its downstream genes, one of which was *PMEI8* (Figs 3C,3D, and 4A-C). Taken together, by integrating both UPB1 and MYB50 transcriptional networks, UPB1 not only controls ROS homeostasis but also cell wall status, such as pectin modification via *PMEI8* which is a direct target of MYB50. However, *PMEI8* regulation alone does not fully elucidate the regulatory mechanisms of root growth in the UPB1 network. Since we also found several transcription factors in the MYB50 direct target genes, analyzing the function of these transcription factors would reveal more details of root growth regulatory mechanisms in the UPB1 gene regulatory network.

**Fig 6.**
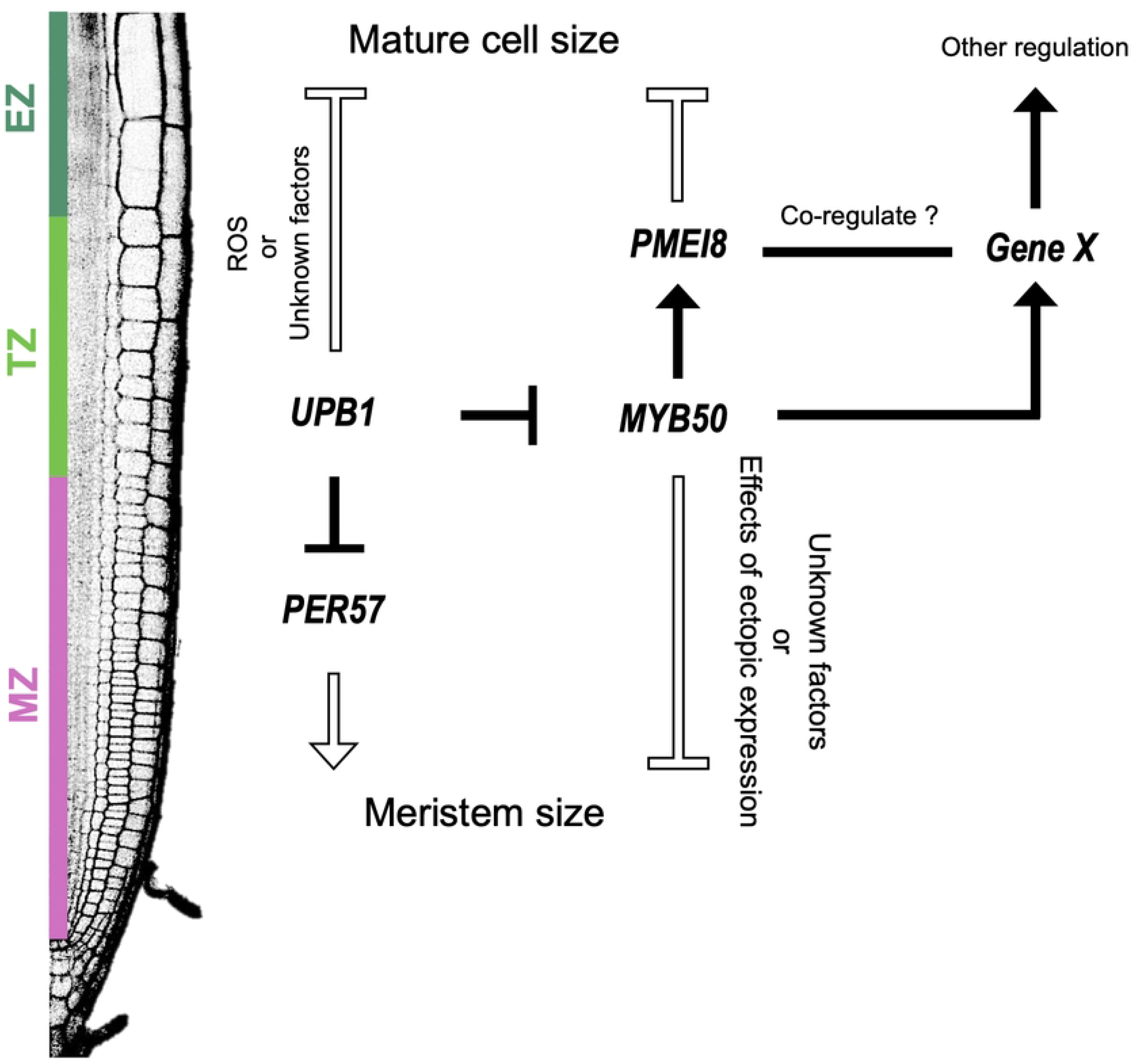
A schematic model of gene regulatory network under UPB1 and MYB50 controls. Black arrows and “T”s indicate transcriptionally direct target genes. The black frame arrow and “T”s indicate an effect on the root phenotype. (MZ): Meristematic zone. (TZ): Transition zone. (EZ): Elongation zone. UPB1 directly represses *PER57* in the TZ and negatively regulates the meristem size [15]. In the EZ, UPB1 directly represses *MYB50* expression. The root phenotypic analysis using *pXVE::YFP-MYB50*/Col-0 revealed that MYB50 negatively regulates the meristem size through a pathway separate from ROS regulation whereas the cause remains unknown. However, *MYB50* was expressed in the EZ close to the differentiation zone and directly activated *PMEI8* expression. *MYB50* regulatory network and *PMEI8* inhibit cell elongation since *pXVE::YFP-MYB50*/Col-0 and *pXVE::YFP-PMEI8*/Col-0 shortened mature cell length. The MYB50- regulated transcriptional network includes several cell wall-associated genes, which may co-regulate with PMEI8 to exert some regulation.

## Acknowledgments

We thank the *Arabidopsis* Biological Resource Center for providing the seeds.

## Data availability

RNA-Seq and ChIP-Seq data were deposited in the DNA Data Bank of Japan (DDBJ) database under the accession codes DRA016078 and DRA016088, respectively.

## Author Contributions

H.T. conceived the study. A. M. and H. T. designed the study. K.M., H.M., S.S., T.T., S.K., and H.T. performed plant genotyping, plasmid construction, plant transformation, and RNA preparation. K.M., S. K., T. T., S. S., and H.T. performed RT-qPCR. T.T., S.K., and H.T. prepared the RNA-Seq library. T.S. performed the RNA sequencing. K.M., T.S., and H.T. analyzed the RNA-Seq data. N.N., S.S. and H.T. performed ChIP and ChIP-Seq library preparation. N.N. performed the ChIP-sequencing. K.M., N.N., T.K., and H.T. analyzed the ChIP-Seq data. M.K. and H.T. performed confocal microscopy and time-lapse imaging. K.M. analyzed the intensity of fluorescent proteins. K.M. and H.T. wrote the manuscript and produced the figures.

## Supporting information

**S1 Fig. Quantification of *MYB50* repression by UPB1 at the single cell level using multi-color reporter line.** Time-lapse images of z-stacks at the early differentiation zone of *pXVE::CFP-cUPB1*/*pMYB50::gMYB50-GFP/upb1-1* were taken by LAS X every 20 min for 5 h. Among 50 z-stacks for each time point, the image which captured the nucleus most clearly was selected because the nucleus moves rapidly in the cells during imaging. CFP and GFP fluorescence of each nuclear signal in the selected images were quantified as a mean intensity by Fiji software. “Z” indicates the z-stack number. “T” indicates the time point after starting time-lapse imaging.

**S2 Fig. *UPB1* and *MYB50* expression in the multicolor reporter line.** *UPB1* and *MYB50* expression in *pXVE::CFP-cUPB1*/*pMYB50::gMYB50-GFP*/*upb1-1* after 5 h of 5 μM estradiol (Est) treatment as measured by RT-qPCR (n = 5, mean ± SD). Significant differences were determined by Student’s *t*-test and compared with untreated plants (*** *p* < 0.001; * *p* < 0.05). *p* value: *UPB1* expression, *p* < 0.001; *MYB50* expression, *p* = 0.0168.

**S3 Fig. Effects of H_2_O_2_ in *MYB50* induced-overexpressor (*pXVE::YFP-MYB50*/Col-0).** (A) Confocal images of the meristematic zone in Col-0 and *pXVE::YFP-MYB50*/Col-0 treated with 5 μM estradiol (Est) or both 5 μM Est and 500 μM H_2_O_2_ treatment for 24 h. Roots were stained with propidium iodide (PI). White arrowheads indicate the ends of the meristematic zone. The blue arrowheads indicate quiescent cells (QC). Scale bar, 100 µm. (B) Quantified cortex cell number in the meristematic zone in Col-0 and *pXVE::YFP-MYB50*/Col-0 upon 5 μM Est or both 5 μM Est and 500 μM H_2_O_2_ for 24 h. Counted root numbers: Col-0 Est = 44, Col-0 Est+ H_2_O_2_ = 38, *MYB50* induced-overexpressor #1 Est = 53, #1 Est+ H_2_O_2_ = 46, #2 Est = 38, and #2 Est+ H_2_O_2_ = 37. Significant differences were determined by Student’s *t*-test compared to Est alone and both 5 μM Est and 500 μM H_2_O_2_ treatment (*** *p* < 0.001).

**S4 Fig. Expression of eight genes that were direct targets of MYB50 and negatively regulated by UPB1.** Expression patterns of selected eight genes from the integration of MYB50 and UPB1 networks in a time-course RNAseq analysis of *pXVE::YFP-MYB50*/Col-0 roots treated with 5 μM Est for 1, 3, or 6 h. The expression values at each time point are relative to the value at 0 h from the RNA-seq results. Error bars indicate SE; (* *q*-value < 0.05; ** *q*-value < 0.01; *** *q*-value < 0.001). *p* value: ERD10 expression after 3 h, *p* = 0.0247; COR47 expression after 3 h, *p* = 0.0045.

**S5 Fig. *PMEI8* expression in the root.** *PMEI8* expression data from root map datasets [32]. Sections 1–6 correspond to the meristematic zone (MZ). Sections 7 and 8 correspond to the elongation zone (EZ). Sections 9–12 describe the differentiation zones (DZ). The tissue (x-axis) represents the radial cell type of the root. Colors indicate *PMEI8* expression levels (color code: YlOrRd; low: red; yellow: white).

**S1 Table.** Primers used in this study

**S2 Table.** Significantly upregulated genes in the time-course RNA-seq used *MYB50* induced-overexpressor

**S3 Table.** Significantly downregulated genes in the time-course RNA-seq used *MYB50* induced-overexpressor

**S4 Table.** List of overlapping genes between MYB50- and UPB1-regulated genes, related to **Fig 4D Supplemental dataset 1.** List of genes in which MYB50 bound 1,000 bp upstream in the ChIP-seq

**S1 Movie. Time-lapse imaging (tile-scan) of *pXVE::CFP-cUPB1*/*pMYB50::gMYB50-GFP*/*upb1-1*.** 6-day-old seedlings of *pXVE::CFP-cUPB1*/*pMYB50::gMYB50-GFP/upb1-1* were transferred from the MS medium to chambered cover glass in media containing 5 µM estradiol. The roots were then stained with PI. Time-lapse images were captured using LAS X every 40 min for 7 h and 20 min.

